# LET-dependence of radiation-induced makers of Immunogenic Cell Death in human cancer cell lines

**DOI:** 10.1101/2022.01.25.477729

**Authors:** Brian Ponnaiya, Anthony LoMastro, Peter W. Grabham, Guy Garty, Andrew D. Harken, Sally A. Amundson, Elizabeth M.C. Hillman, David J. Brenner

## Abstract

**Purpose:** It has been suggested that heavy-ion radiation therapy may contribute to the control of distal metastases. These distant responses may include immune cell activation. Immunostimulation resulting from radiation-induced immunogenic cell death (ICD) of cancer cells, leads to the recruitment of anti-tumor T cells. Specific markers of ICD include translocation of calreticulin (CRT) and extracellular release of high mobility group box 1 protein (HMGB1), and ATP. However, the LET dependence of these effects remains unknown.

**Materials and Methods:** Expression of the molecular indicators described above were tested in a panel of human cancer cell lines, that included pancreatic cancer (Panc1 and Paca2), glioblastoma (U87 and LN18) and melanoma (HTB129 and SK-Mel5). Cells were irradiated with 5 Gy of particles spanning a range of LETs, from 10 KeV/μm to 150 KeV/μm and assayed for relocalization of calreticulin and release of HMGB1 and ATP were assayed 24 hours later.

**Results:** In the pancreatic cancer cell lines (Panc1 and Paca2) there was a continued increase in the membrane relocalization of calreticulin as a function of increasing LET up to 150 KeV/μm. The melanoma cell lines, HTB129 and Sk-Mel5 showed similar patterns. In contrast, calreticulin levels were higher, but not LET-dependent, in irradiated U87 and LN18 (glioblastoma) lines. With the exception of the response in Paca2, increases in LET correlated with increases in HMGB1 that seemed to peak at 100 KeV/μm and then either remain unchanged or decrease at 150 KeV/μm. while the ATP levels were elevated in media from some of the irradiated groups, there were no clear patterns either by cell type or LET.

**Conclusions:** Our results indicate that at equal doses, although there is an overall trend of increases in the responses to increasing LETs, there are significant cell line-specific differences in the patterns of expression of these key ICD markers.

## INTRODUCTION

There is now a growing amount of evidence that heavy ion beam radiotherapy provides better local control of various types of primary cancers, including pancreatic, GI and hepatocellular cancers, when compared to conventional photon irradiation (eg (Schulz-Ertner, Nikoghosyan et al. 2007, Abousaida, Seneviratne et al. 2021, Liermann, Ben-Josef et al. 2021, Noticewala and Das 2021, Yamada, Takiyama et al. 2021). Significantly, heavy ion therapy also appears to show high efficacy at controlling distal metastases and it is thought that particle irradiation may suppress the metastatic potential of cancer by modifying the host antitumor immunity (Ogata, Teshima et al. 2005, Matsunaga, Ueda et al. 2010, Durante, Brenner et al. 2016).

However, there are two central questions that remain unanswered. The first is what is the optimal heavy ion or LET to trigger such a response? Heavy ion therapies conducted in Europe and Japan have almost exclusively used carbon ions where the LET range in the spread-out Bragg peak is between ~50-200 KeV/um (Kanematsu, Matsufuji et al. 2018). However, there has not been a systematic evaluation of the efficacy other LETs / ions that might be technically easier and significantly cheaper to produce and deliver, for example, a helium ion beam with a spread-out Bragg peak between (~8-50 keV/μm) (Kempe, Gudowska et al. 2007). To conduct mono-LET in vitro studies, we have developed a range of mono-LET ion beams using our 0-2.5 MeV/amu Singeltron accelerator and linear accelerator booster (the latter providing beams at 4.1 and 5.5 MeV/amu) (Marino, 2017). These beams maintain near constant LET over a depth suitable for irradiating cells in culture (typical thickness: ~5 μm), with available LETs from 10 to 150 keV/μm, spanning the range of LET values relevant to HIRT.

The second central question of HIRT is what are the mechanistic bases for why heavy ions are so effective in controlling distal secondary tumors? One hypothesis is that heavy ion therapy might work through the immune system via the immunogenic cell death pathway. Immunogenic cell death is a functionally unique form of stress-driven regulated cell death that results in the activation of cytotoxic T lymphocyte (CTL)-driven adaptive immunity as well as long-term immune memory (Zhou, Wang et al. 2019) This specific type of cell death has been observed in response to microbial pathogens as well as chemo- and radio-therapies (Obeid, Tesniere et al. 2007, Fucikova, Kralikova et al. 2011, Golden, Pellicciotta et al. 2012, Golden, Frances et al. 2014, Campisi, Barbet et al. 2016). Cells dying by this mechanism exhibit three distinct cellular responses; cell surface translocation of calreticulin (CRT), and extracellular release of both high-mobility group box 1 (HMGB1) and ATP (Galluzzi, Vitale et al. 2020). These three markers have also been used to identify ICD in both in vivo and *in vitro* systems.

Here we present work that using the mono-LET beams available at RARAF with a range of 10 to 150 KeV to examine the LET dependence of radiation-induced expression of the three markers of ICD in a panel human cancer cell lines.

## MATERIALS AND METHODS

### Cell culture and Irradiation

Human pancreatic cell lines, Panc1 and Paca2, melanoma lines Sk-Mel5 and HTB129, and glioblastoma lines U87 and LN18 were obtained from ATCC. Cells were maintained in either Eagles or Dulbecco’s media supplemented with 10% FBS as recommended.

Irradiations were conducted as previously described ((Geard, Jenkins-Baker et al. 2002)). Briefly cells were seeded on 6 μm mylar dishes at a concentration of 2X10^5^ per dish (2 ml media per dish) and 24 hours later were exposed to 5 Gy of particles with a range of LETs (10-150 keV/μm). Irradiated dishes were fixed with formalin at 24 hours post irradiation.

### Irradiation Set up and dosimetry

Irradiations were performed at the Radiological Research Accelerator Facility’s (RARAF) Track Segment “mono-LET” irradiation platform (Bird, Rohrig et al. 1980, Marino 2017), which provides irradiation of thin biological samples using proton, deuteron and helium beams. Briefly, an ion beam from the RARAF accelerator is energy selected, using a set of magnetic dipoles, defocused and exits the vacuum system through a 2.9 μm thick Havar window. To mimic a broad beam irradiation cell monolayers in a 30 mm diameter holder with 6 μm mylar bottom are scanned across the beam. Assuming a thin target, this results in a “segment” of a particle track, having nearly constant LET, traversing the sample.

Dosimetry is performed prior to each irradiation, using a set of detectors irradiated in the same geometry as the cell monolayers (Garty 2022). A set of wipe-off vanes are used to monitor beam current and control the scanning speed of samples and detectors across the window, to compensate for variations in dose rate. A unit gain tissue equivalent proportional counter (TEPC), operated in pulse mode is used to measure the linear energy transfer (LET) of the ions. A second TEPC, operated in current mode is used to measure dose and calibrate the monitor vanes, prior to each irradiation. A lithium drifted silicon solid state detector is used to verify beam energy and measure uniformity across the sample irradiated.

### Immunocytostaining and Image Analysis

Formalin fixed cells were blocked with 2% BSA in PBS for 30 minutes at room temperature. Cells were incubated with the anti-Calreticulin antibody, ab2907 (1:75), for 1 hour at room temperature, washed with PBS, and incubated with the secondary antibody ab150077 Alexa Fluor® 488 goat anti-rabbit IgG (H+L) used at 2μg/ml for 1h. DAPI was used to stain the cell nuclei (blue) at a concentration of 1.43μM. Images were taken at 20X magnification and analyzed for mean cellular fluorescence using Image ProPlus. Cellular intensities from 100-150 cells per group, per experiment, were averaged, and the means of three separate experiments were calculated.

### Media Analysis

24 hours post irradiation, 250 μl media were removed from individual dishes and either assayed immediately for HMGB1 or ATP levels, or stored at −80°C for future analysis. Levels of HMGB1 in culture media were assayed using a sandwich ELIZA kit (LifeSpan BioSciences) following the manufacturers protocols. ATP levels were measured using the luciferase based ENLITEN® ATP Assay System (Promega).

### Statistical Analysis

Data from three independent experiments were calculated as means and either standard error of means for calreticulin or standard deviations for HMGB1 and ATP. Significant differences among different groups were determined by unpaired Student’s t-test with a 2-tailed distribution. P values of <0.05 was considered statistically significant.

## RESULTS

The LET-dependent responses of ICD markers were assessed in six human cell lines representing 3 common cancer types that are candidates for HIRT: pancreatic cancer (Panc1, PaCa-2), glioblastoma (U87, LN18) and melanoma (SK-Mel-5, HTB129). Cells were exposed to 5 Gy of particles spanning a range of LETs (10, 65, 100 and 150 KeV/μm) and relocation of calreticulin, and release of HMGB1 and ATP were assayed 24 hours post-irradiation.

### Expression of Calreticulin on cell membrane

In the pancreatic cancer lines, Panc1 and Paca2, background levels remained between 500-550 and 550-700 AU respectively (Fig 2A, 2B). In response to radiation, levels in Panc1 cells increased to around 800 AU at 10 KeV/μm with similar levels at 65 KeV/μm. Levels were further increased to 1050 AU at 100 and 150 KeV/μm. Calreticulin intensities in Paca2 cells irradiated with 10 KeV/μm were 900 AU, with a plateau at about 1100 AU at 65 and 100 KeV/μm and a further increase to 1700 AU at 150 KeV/μm.

**FIGURE 1:**
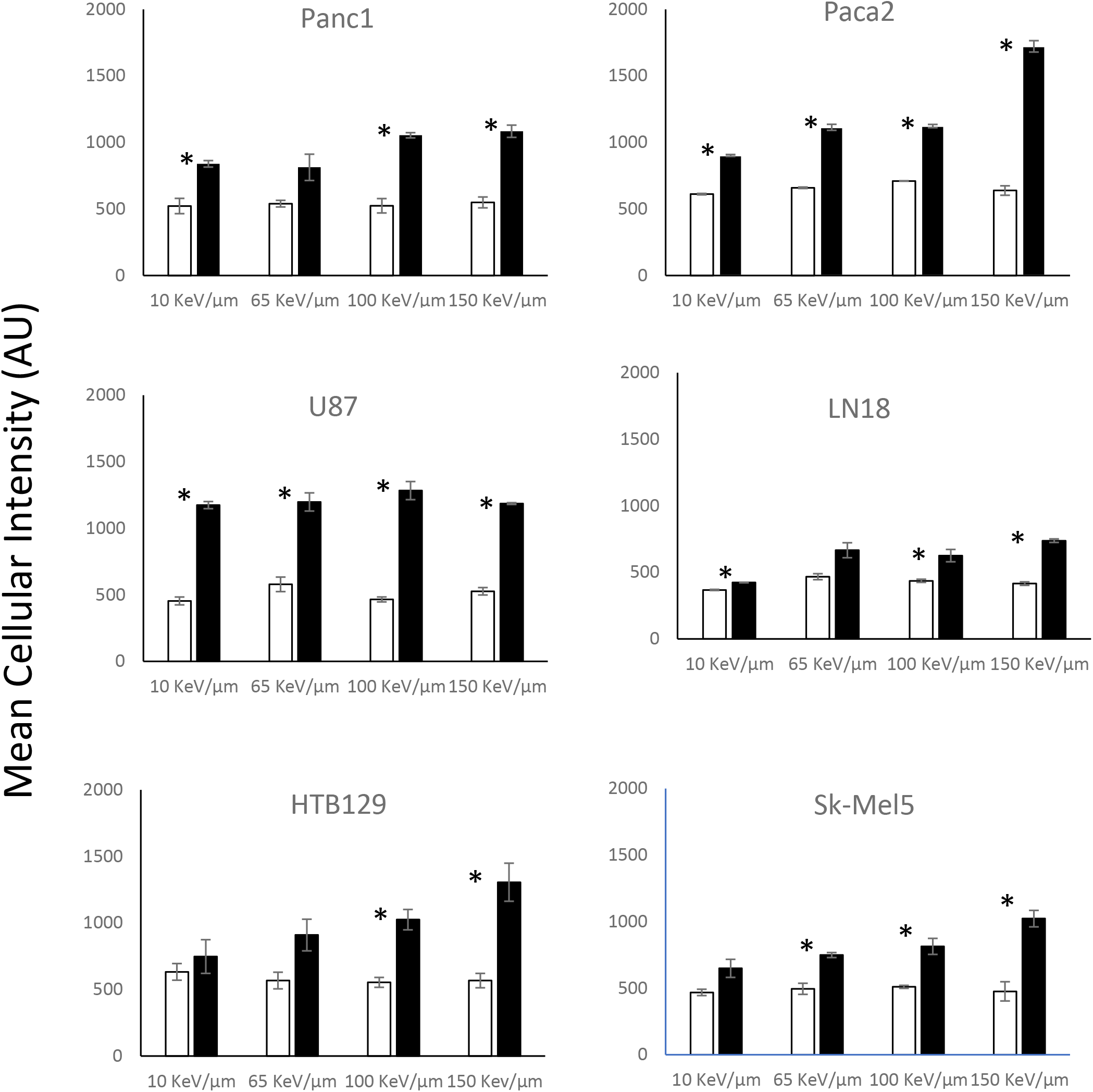
Cell membrane expression of calreticulin in human cancer cell lines in control (clear bars) and 24 hours post exposure to 5 Gy of ions at the indicated LET (filled bars) (mean ± SEM; * represents P<0.05 between irradiated and corresponding control groups)

**FIGURE 2:**
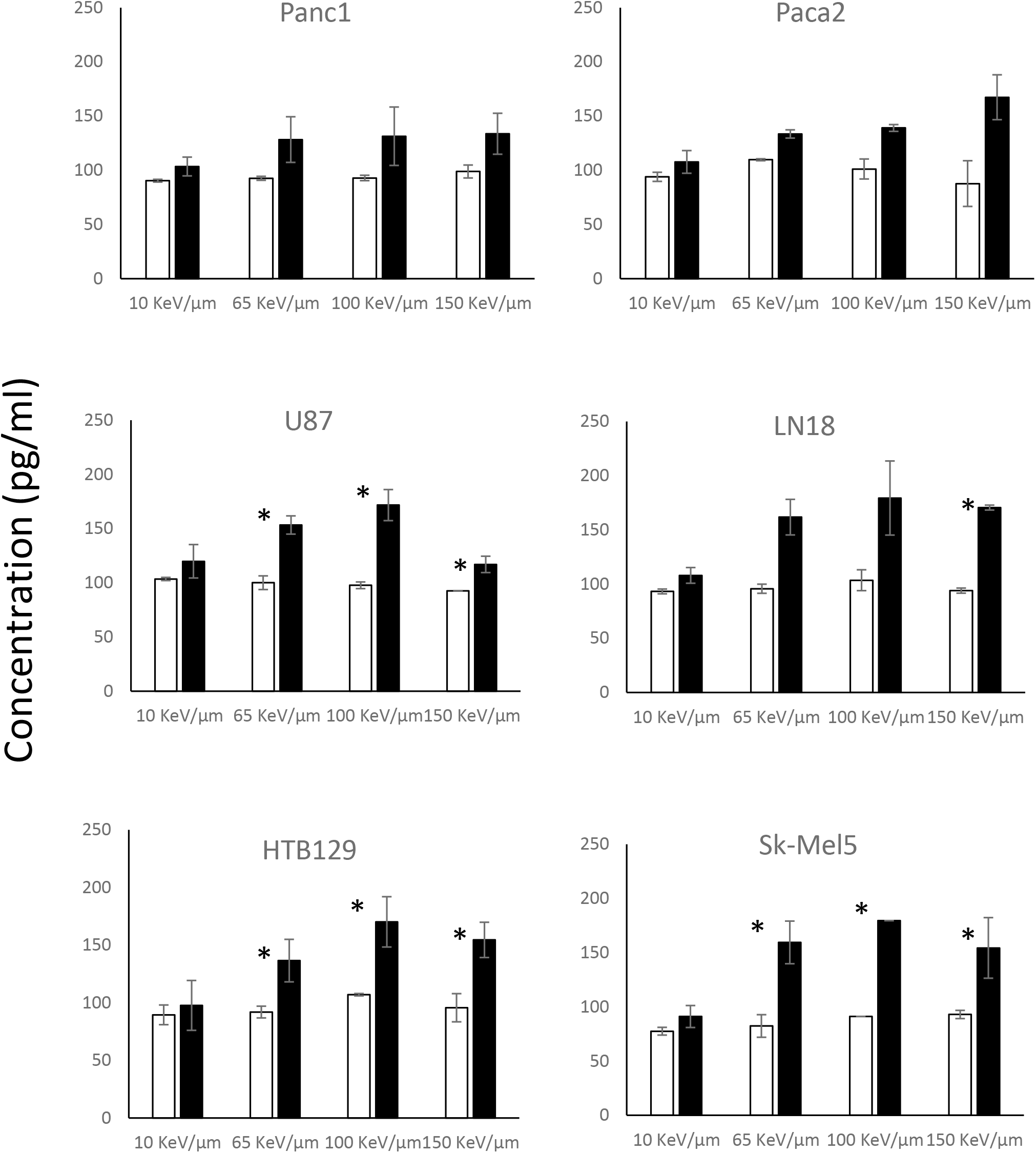
Detection of HMGB1 in the cell media in control (clear bars) and irradiated dishes (filled bars) 24 hours post exposure to 5 Gy of ions at the indicated LET (mean ± SD; * represents P<0.05 between irradiated and corresponding control groups).

**FIGURE 3:**
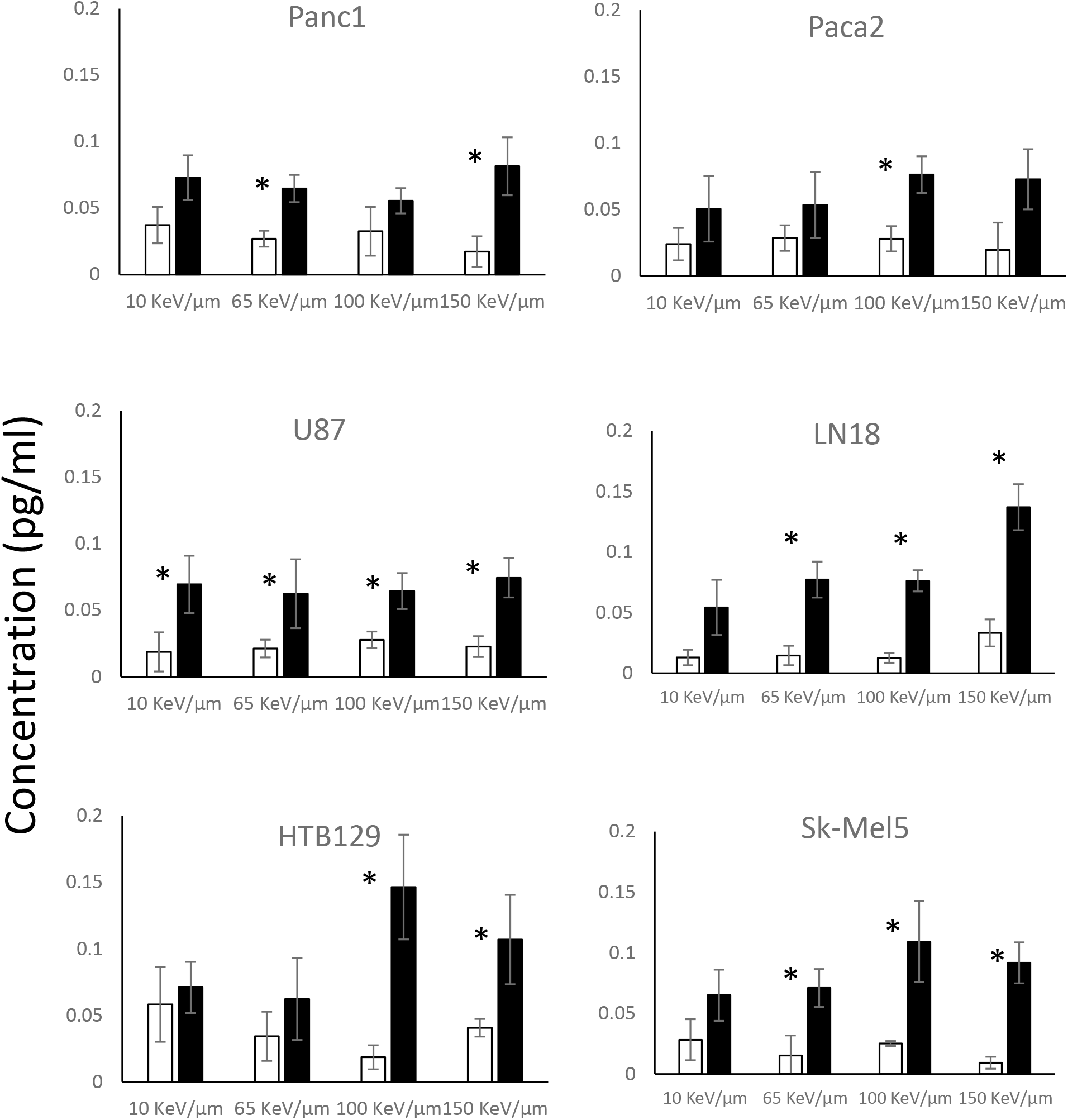
Detection of ATP in cell media in control (clear bars) and irradiated dishes (filled bars) 24 hours post exposure to 5 Gy of ions at the indicated LET (mean ± SD; * represents P<0.05 between irradiated and corresponding control groups).

In the melanoma cell lines, control levels were between 470-500 AU in Sk-Mel5 and 550-630 AU in HTB129 (Fig 2C, 2D). In both lines, there were small, but consistent, increases as a function of increasing LET. In Sk-Mel5, CRT intensities went from 650 AU at 10 KeV/μm to 1020 AU at 150 KeV/μm. Similarly, in the HTB129 line, intensities increased from 750 AU at 10 KeV/μm to 1300 AU at 150 KeV/μm.

In the glioblastoma lines, background intensities were 370-470 AU in LN18 and 450-580 AU in U87 (Fig 2E, 2F). With the exception of LN18 at 10 KeV/μm, both glioblastoma lines had similar responses to radiation, increasing to 1100-1200 AU at all LETs in the U87 line and 600-700 AU in the LN18 line.

### Extracellular HMGB1

In the pancreatic cell lines, levels of HMGB1 detected in the media at 24 hours post irradiation was between 90-100 pg/ml in unirradiated controls. In the irradiated Panc1 groups, HMGB1 levels ranged from 100 pg/ml at 10 KeV/μm to 130 pg/ml at 65,100 and 150 KeV/μm, representing a 1.3-1.4 fold increase over the corresponding controls. In the Paca2 samples, again, levels at 10 KeV/μm were similar to controls, but were elevated at higher LETs, ranging from 135 pg/ml at 65 and 100 KeV/μm up to 169 pg/ml at 150 KeV/μm.

In the glioblastoma lines, 10 KeV/μm induced little to no increase in HMGB1 levels when compared to controls. At higher LETs there was a 1.5 fold increase up to 100 KeV/μm. Interestingly, there was no further increase at 150 KeV/μm and further the media from U87 cells irradiated with 150 KeV/μm particles seemed to have lower levels of HMGB1 than that at 65 or 100 KeV/μm.

Both irradiated melanoma cells (SK-Mel5 and HTB129) released increasingly higher LET dependent levels of HMGB1 into the media when compared to corresponding control cultures. These increases ranged from 135-155 pg/ml for the HTB129 cells, and 155-180 pg/ml in the Sk-Mel5 cultures (representing 1.5 and 1.6-2 fold increases over control cultures respectively). At 150 KeV/μm however, there was a slight decrease in the response.

### Extracellular ATP

ATP levels in media from nonirradiated pancreatic cell cultures ranged between 0.02-0.04 pg/ml. 24 hours post-irradiation, ATP levels increase to 0.05-0.08 pg/ml in the media from irradiated Panc1 cultures, and 0.05-0.076 pg/ml in the Paca2 cultures.

In the glioblastoma cell lines, ATP levels in media from nonirradiated cultures ranged between 0.01-0.03 pg/ml. Following irradiation these levels increased to 0.063-0.075 pg/ml in the media from irradiated U87 cultures, and 0.05-0.14 pg/ml in the LN18 cultures. Media from control Sk-Mel 5 and HTB129 cultures had 0.01-0.03 pg/ml and 0.02-0.05 pg/ml ATP respectively. In response to irradiation, these levels increased to 0.06-0.15 pg/ml in the irradiated HTB129 groups and 0.065-0.1 pg/ml in the Sk-Mel5 cultures.

## DISCUSSION

The current study focused on the radiation-induced expression of ICD markers as a function of increasing LETs in human cell lines representing three different cancer types. From the data presented here, there appears to be cell-type specific differences in the membrane relocalization of calreticulin as a function of LET. Both pancreatic cancer cell lines (Panc1 and Paca2) demonstrated a continued increase in the membrane expression as a function of increasing LET up to 150 KeV/μm. Similar responses, albeit to a lesser degree, were observed in the melanoma lines, HTB129 and Sk-Mel5. In contrast, irradiation of the glioblastoma cell lines (U87 and LN18) resulted in elevated levels of calreticulin that was not LET dependent; while almost all irradiated populations had increased membrane calreticulin when compared to controls, there were no significant differences in levels between any of the irradiated populations at different LETs. Release of HMGB1 showed a somewhat different pattern. Except for the Paca2 response, increases in LET correlated with increases in HMGB1 that seemed to peak at 100 KeV/μm and then either remain unchanged or decrease at 150 KeV/μm. In support, this peak in response at 100 KeV/μm has been previously described and attributed to the fact that the average separation of ionizing lesions at this particular LET coincide with the diameter of cellular DNA (Hall and Giaccia 2006). In fact studies conducted at RARAF using the mono-LET beams used here, demonstrated that at equal doses, chromosomal aberrations in chinese hamster V79 cells (induced by DNA double strand breaks) increased as a function of increasing LET up to about 100 KeV/μm and then decreased (Geard 1985). Furthermore, the link between DNA double strand breaks and high-LET induced ICD has previously been made by Durante and Formenti (Durante and Formenti 2018) who proposed that following high-LET radiation, a high fraction of double stranded DNA fragments are produced that can be released in the cytoplasm following nuclear envelope breakdown, triggering the mechanisms underlying immunogenic cell death.

Radiation-induced ICD in *in vitro* systems has been well documented for x-rays and more recently for high-LET ions. A recent report compared the relocalization of calreticulin as a function of exposure to either photon, proton (~2 KeV/μm) or carbon ion (~30 KeV/μm) beams (Huang, Dong et al. 2019). Protons and photons at similar doses produced the same levels of increases in calreticulin, and carbon ions induced higher levels at 2 and 4 Gy but not at 10 Gy. As in the present study, they found that increases in membrane calreticulin were tumor-type dependent. In cells from radiation-sensitive tumors, such as nasopharyngeal carcinoma (CNE-2) and tongue squamous carcinoma (Tca-8113), carbon ions induced higher levels of membrane calreticulin as compared to photons or protons at both 2 and 4 Gy. However, this was not the case in a glioblastoma line, U251, where exposure to carbon ions did not induce significantly different levels of calreticulin when compared to photons or protons at these doses. Similarly, the release of HMGB1 as a function of exposure to carbon ions at two different LETs demonstrated that exposure to 70 KeV/μm ions induced higher levels of HMGB1 release when compared to that resulting from irradiation with 13 KeV/μm at equilethal doses (Onishi, Okonogi et al. 2018) In conclusion, we have demonstrated the radiation-response of key markers of ICD in cell lines representing three specific cancer types that might benefit from HIRT. The data suggests that although there are LET-dependent increases in many cases, there are also large variations between the types of cancer, and even between individual cancers of the same type.

## ACKNOWLEDGEMENTS

The studies reported here were supported by U01CA236554 from NCI and U19AI067773 from NIAID.

## BIOGRAPHICAL NOTE

**Brian Ponnaiya** is a Research Scientist at the Center for Radiological Research at Columbia University, NY.

**Anthony Lomastro** was a Technician based at the Radiological Research Accelerator Facility (RARAF) that is part of the Center for Radiological Research at Columbia University, NY.

**Guy Garty** is an Associate Professor of Radiation Oncology (in the Center for Radiological Research) at Columbia University and the Associate Director of RARAF.

**Andrew D. Harken** is an Associate Research Scientist at RARAF and the Center for Radiological Research at Columbia University.

**Peter W. Grabham** is an Assistant Professor of Radiation Oncology (in the Center for Radiological Research) at Columbia University

**Sally A. Amundson** is an Associate Professor of Radiation Oncology (in the Center for Radiological Research) at Columbia University

**Elizabeth M.C. Hillman** is the Herbert and Florence Irving Professor at the Zuckerman Institute at Columbia University and a Professor of Biomedical Engineering and radiology (Physics) at Columbia University.

**David J. Brenner** is the Higgins Professor of Radiation Biophysics (in Radiation Oncology) and of Environmental Health Sciences and is the Director of the Center for Radiological Research (CRR) and the Director of RARAF.

